# AI-powered Deep Visual Proteomics reveals critical molecular transitions in pancreatic cancer precursors

**DOI:** 10.1101/2025.07.07.663528

**Authors:** Jimin Min, Lisa Schweizer, Gijs Zonderland, Benson Chellakkan Selvanesan, Lukas Oldenburg, Seong-Woo Bae, Bong Jun Kim, Benjamin J. Swanson, Kelsey A. Klute, Thomas C. Caffrey, Paul M. Grandgenett, Michael A. Hollingsworth, Ishani Ummat, Maximilian T. Strauss, Andreas Mund, Anirban Maitra

**Author notes:** These authors contributed equally to this work (Jimin Min, Lisa Schweizer). **Correspondence:** Andreas Mund, Ph.D. OmicVision Biosciences, BioInnovation Institute, Ole Maaløes Vej 3, DK 2200 Copenhagen N, Denmark., and Anirban Maitra, MBBS. 1515 Holcombe Blvd Z3.3040, The University of Texas MD Anderson Cancer Center, Houston, TX 77030.

## Abstract

Pancreatic ductal adenocarcinoma (PDAC) evolves through non-invasive precursor lesions, yet its earliest molecular events remain unclear. We established the first spatially resolved proteomic atlas of these lesions using Deep Visual Proteomics (DVP). AI-driven computational pathology classified normal ducts, acinar-ductal metaplasia (ADM), and pancreatic intraepithelial neoplasia (PanIN) from cancer-free organ donors (incidental, “iPanINs”) and PDAC patients (cancer-associated, “cPanINs”). Laser microdissection of 96 discrete regions containing as few as 100 phenotypically matched cells and ultrasensitive mass spectrometry quantified a total of 8,512 proteins from formalin-fixed tissues. Distinct molecular signatures stratifying cPanINs from iPanINs, and remarkably, many cancer-associated proteins already marked histologically normal epithelium. Four core programs - stress adaptation, immune engagement, metabolic reprogramming, mitochondrial dysfunction - emerged early and intensified during progression. By integrating DVP with AI-guided tissue annotation, we demonstrate that molecular reprogramming precedes histological transformation, creating opportunities for earlier detection and interception of a near-uniformly lethal cancer.

**Significance:** Our spatially-resolved proteomics atlas uncovers distinct molecular signatures in pancreatic cancer adjacent precursor lesions, clearly diverging from those in incidental, cancer-free pancreatic lesions. Our deep proteomics dataset offers a valuable resource for identifying novel biomarkers and therapeutic targets, informed by the earliest cancer-associated molecular events in archival pancreatic tissues.

## Introduction

Pancreatic ductal adenocarcinoma (PDAC) remains one of oncology’s most formidable challenges, with a dismal five-year survival rate below 10% owing to late detection, therapeutic resistance, and rapid metastatic spread (1). The canonical progression model of PDAC depicts a stepwise transition from histologically unremarkable normal ductal epithelium through non-invasive precursor lesions known as pancreatic intraepithelial neoplasia (PanIN) to invasive carcinoma, driven by sequential accumulation of oncogenic *KRAS* activation and inactivation of *CDKN2A*, *TP53* and *SMAD4* (2,3). Acinar-to-ductal metaplasia (ADM) are “bi-phenotypic” lesions with features of both acinar and ductal cells and are considered an intermediate transitional step between normal ductal epithelium and PanINs, serving as “precursors to the precursor” (4). In contrast to PanIN lesions, the biology of ADMs in humans is poorly described and much of the knowledge stems from studies in murine models (5).

Recent studies in cancer-free organ donor samples have demonstrated a remarkably high prevalence of incidental PanIN lesions (∼60% of the general population), including in young adults (6). Given the relatively uncommon incidence of PDAC, most incidental PanIN (“iPanINs”) will not progress to invasive neoplasia during an individual lifetime. Nonetheless, whether iPanINs harbor a distinct molecular signature from precursor lesions adjacent to PDAC (cancer-associated or “cPanINs”) remains unknown. Bridging this knowledge gap is crucial. Meaningful early detection and cancer interception strategies must identify which subset of precursor lesions are most likely to progress to PDAC while avoiding surveillance and overdiagnosis pitfalls for the rest (7).

Technical challenges compound these knowledge gaps: low-volume liquid biopsies often fail to detect circulating tumor DNA in early-stage disease; bulk omics approaches sacrifice critical spatial context; antibody-based methods offer limited proteomic coverage constraining discovery potential; and mRNA levels do not necessarily correspond to protein abundance. Moreover, given low accessibility of organ donor pancreas samples, nearly all prior studies have predominantly analyzed precursor lesions from patients already diagnosed with PDAC (8,9), potentially missing *de novo* molecular events occurring in lesions from cancer-free individuals. Growing evidence also indicates a more complex biological reality with heterogeneous evolutionary trajectories of precursor lesions profoundly influenced by the precancer microenvironment (PME) (6,9,10). The concept of “field cancerization” (11) − molecular alterations extending beyond visible lesions into surrounding normal-appearing tissues − plays a critical role in PDAC development, potentially driving the high recurrence rates following surgical resection.

We overcome these constraints through Deep Visual Proteomics (DVP) − integrating high-resolution microscopy, AI-powered computational pathology, targeted laser microdissection, and ultrasensitive mass spectrometry to enable comprehensive protein analysis within precisely defined cell populations (12–14). In this study, we create the first spatially resolved proteomic cartography of multistep PDAC development by analyzing precisely defined tissue compartments across the spectrum from normal ducts to non-invasive precursor lesions to invasive carcinoma. We hypothesized that molecular reprogramming might precede histological changes and that cancer context within the contiguous organ could profoundly influence the cellular state of otherwise histologically normal tissue. By comparing normal ducts and PanINs from both cancer-free individuals and from patients with PDAC (“iPanINs” *versus* “cPanINs”, respectively), we identify critical molecular transitions during pancreatic carcinogenesis and establish a framework for risk stratification of precursor lesions. We also generate the first proteomic atlas of human ADMs. This work addresses fundamental questions about the earliest events in PDAC development, with direct implications for transforming clinical practice from late-stage intervention to early detection and targeted prevention strategies (15).

## Results

### Deep Visual Proteomics workflow and dataset overview

To study the spatial and molecular progression of pancreatic ductal adenocarcinoma (PDAC) at its earliest stages, we implemented Deep Visual Proteomics (DVP) (12), an integrated spatial proteomic approach that combines complementary cutting-edge technologies (**Fig. 1A)**. We analyzed 96 discrete histologically annotated and microdissected regions from 15 individuals, including 10 cancer-free organ donors (incidental or “iCohort”) and 5 patients with PDAC (cancer-associated or “cCohort”), capturing the full spectrum from normal ducts to invasive carcinoma. Our DVP workflow uses 3-μm-thick sections of formalin-fixed paraffin-embedded (FFPE) tissues stained with hematoxylin and eosin (H&E). After acquiring high-resolution images, we employed H-optimus-0, a pathology foundation model trained on over 500,000 histopathology slides, to identify phenotypic clusters across the progression spectrum. This AI-powered approach enhanced our ability to precisely distinguish morphologically similar but biologically distinct tissue compartments at the cellular level of resolution.

**Fig. 1.**
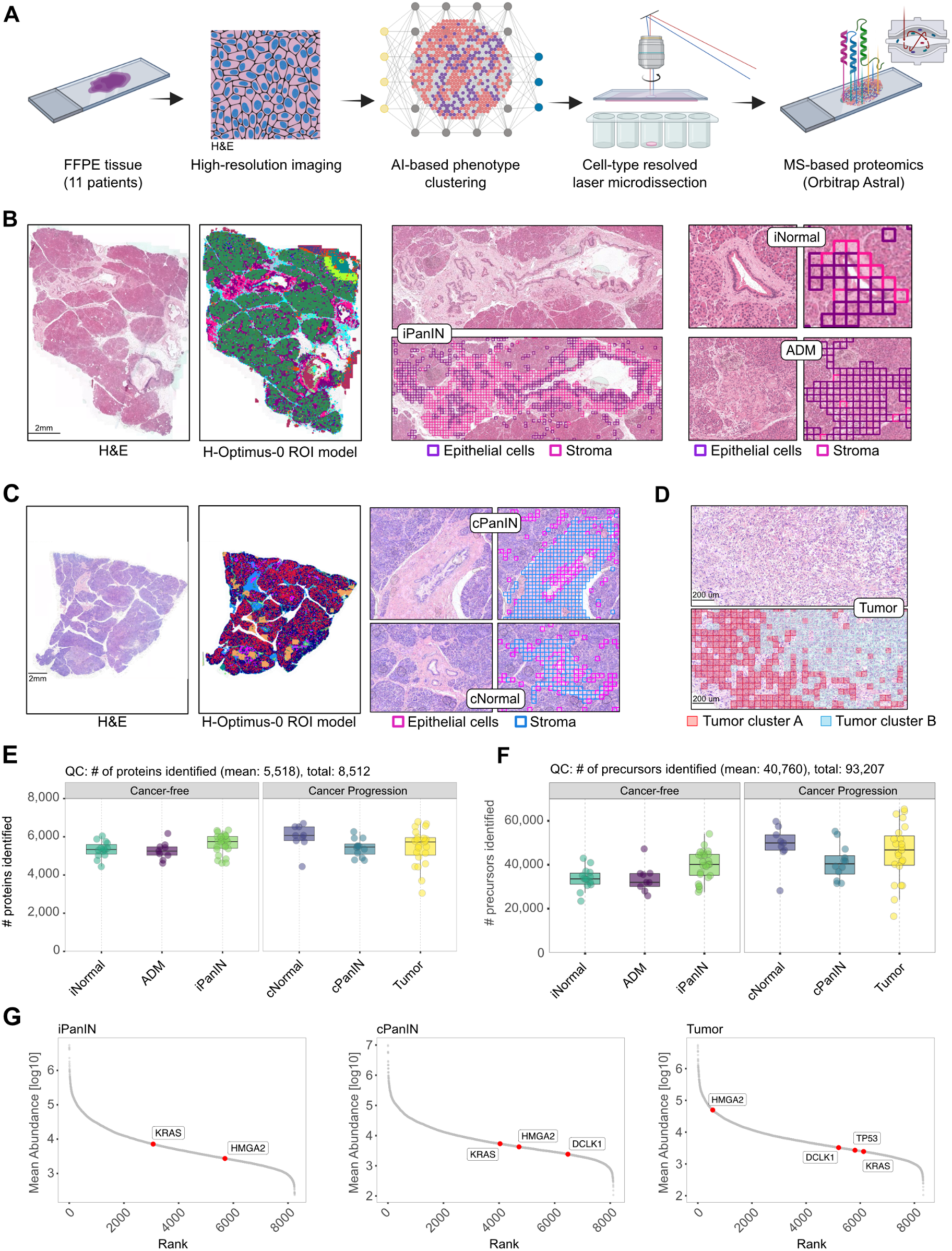
Deep Visual Proteomics of the multistep progression of PDAC. **A)** Schematic overview of the Deep Visual Proteomics (DVP) workflow. FFPE tissue specimen were H&E stained. High-content images were analyzed using AI-based phenotype clustering to extract tissue morphologies and disease-associated pathologies. Cells were extracted using a Leica LMD7 laser-microdissection instrument and cell-type specific proteomes were acquired on an Orbitrap Astral mass spectrometer. **B-D)** Pathology foundation models. H-optimus-0 regions of interest (ROI) model was applied on H&E images to identify iPanIN, iNormal and ADM lesions **(B)**, cPanIN and cNormal **(C)** and representative tumor clusters **(D)**. **E,F)** Number of proteins **(E)** and peptide precursors **(F)** quantified across all cohort groups using MS-based proteomics. **G)** Mean ranked abundance of proteins found in iPanIN, cPanIN, and tumor groups. The abundance of key markers of PanINs or PDAC are highlighted in red (KRAS, HMGA2 DCLK1, TP53). Abbreviations – iPanIN and iNormal = incidental PanIN and adjacent normal ductal epithelium from cancer-free organ donors; cPanIN and cNormal = cancer-associated PanIN and normal ductal epithelium from pancreas with PDAC.

Guided by the pathology foundation model and pathologist annotation, we selected six biological groups for proteomic profiling: normal ducts (“iNormal”, “cNormal”), precursor lesions (ADM, “iPanIN”, “cPanIN”), and invasive carcinoma (PDAC) (**Fig. 1B-D**, **Fig. S1A, S1B**). Using laser microdissection with high spatial precision, we isolated small regions containing approximately 100 phenotypically matched cells for each sample. This targeted approach enabled us to overcome the limitations of bulk tissue analysis, which can mask critical signals from rare cell populations and dilute compartment-specific molecular signatures. We performed subsequent proteomic profiling on the Orbitrap Astral mass spectrometer, quantifying 8,512 unique proteins (**Fig. 1E**) and 93,207 unique peptide precursors (**Fig. 1F**) across all tissue states. The proteins spanned six orders of magnitude in abundance, demonstrating the exceptional sensitivity and dynamic range of our approach (**Fig. 1G, Table S1**). Our quality control analyses confirmed the presence of peptides corresponding to pivotal PDAC-associated genomic drivers across the proteomic abundance range, including KRAS and TP53, established cancer cell markers like HMGA2 (16), and stem cell markers like DCLK1 (17) (**Fig. 1G, S2A**). We reliably identified canonical PanIN markers, including ANXA10, S100P, CLDN18 and MUC5AC, at expected levels across PanIN lesions regardless of cancer context (**Fig. S2B**. Furthermore, when we compared our human atlas on PDAC precursor lesions to recent work in mice (14), we found a substantially higher number of identified proteins as well as a unique set of more than 6,000 proteins defined specifically for human patients (**Fig. S2C**). These findings confirmed our ability to detect known pancreatic neoplasia-associated proteins and the clinical relevance of our proteomic atlas. This integrated methodology allowed us to analyze thousands of proteins within spatially and phenotypically defined cell populations, directly linking molecular profiles to cellular identity and tissue context across the multistep progression of human pancreatic neoplasia.

### Cancer-specific molecular signatures differentiate PDAC from precancers

Our analysis of total data variance across all regions, including both tumor and non-tumor compartments, revealed a clear, not unexpected, separation of the PDAC proteome from normal ducts, ADM, and PanINs (**Fig. 2A**). Differential expression and pathway enrichment analysis identified tumor-specific activation of cancer-related signatures, including extracellular matrix (ECM) remodeling, MAPK, BRAF, and MET signaling pathways, along with significant enrichment of proteins associated with chemoresistance (e.g. ABCG2, ALDH1A1) and tumor-stromal crosstalk (e.g. POSTN, S100A4), underscoring the functional and biological relevance of our proteomic dataset (**Tables S2, S3**). Notably, key contributors included LGALS1 (Galectin-1), a well-established regulator of tumor-stroma interactions (18,19), and SERPINH1 (HSP47), an ECM-associated chaperone involved in collagen processing (20,21) (**Fig. 2B**). Intriguingly, both PLOD3 and COLGALT1, which form a dimeric complex involved in collagen post-translational modifications (PTMs), known as the KOGG complex (22), were upregulated specifically in tumor compartments, suggesting a critical role in ECM remodeling during tumor progression (**Fig. 2B**). Tumor samples displayed molecular hallmarks of cancer consistent with late-stage progression. Elevated TP53 levels appeared specifically in tumors, likely representing stabilization of mutant protein (**Fig. S3A**). DCLK1, a putative cancer stem cell marker, showed progressive increases from cPanIN to tumor (**Fig. S3B**). We identified a distinct set of proteins exclusively expressed in tumor samples but absent in precursor lesions (**Fig. S3C, Table S4**). These tumor-specific proteins participate in key cancer-associated processes including cell cycle regulation (CCNB1, CDC20), DNA replication (GINS1), transcriptional control (RUNX2, FOSL1, GJA1), ECM interaction (FBN2), immune modulation (IFITM3), and polyamine metabolism (SMOX). Although high-grade PanIN lesions were not included in our series, the exclusive appearance of these factors at the PDAC stage likely reflects the acquisition of cell autonomous proliferative and invasive capabilities. This final transformation to invasive carcinoma likely represents the culmination of the extended evolutionary timeline estimated in genetic studies, where a median time of several years from the most recent common ancestor to clinically apparent PDAC has been proposed (23).

**Fig. 2.**
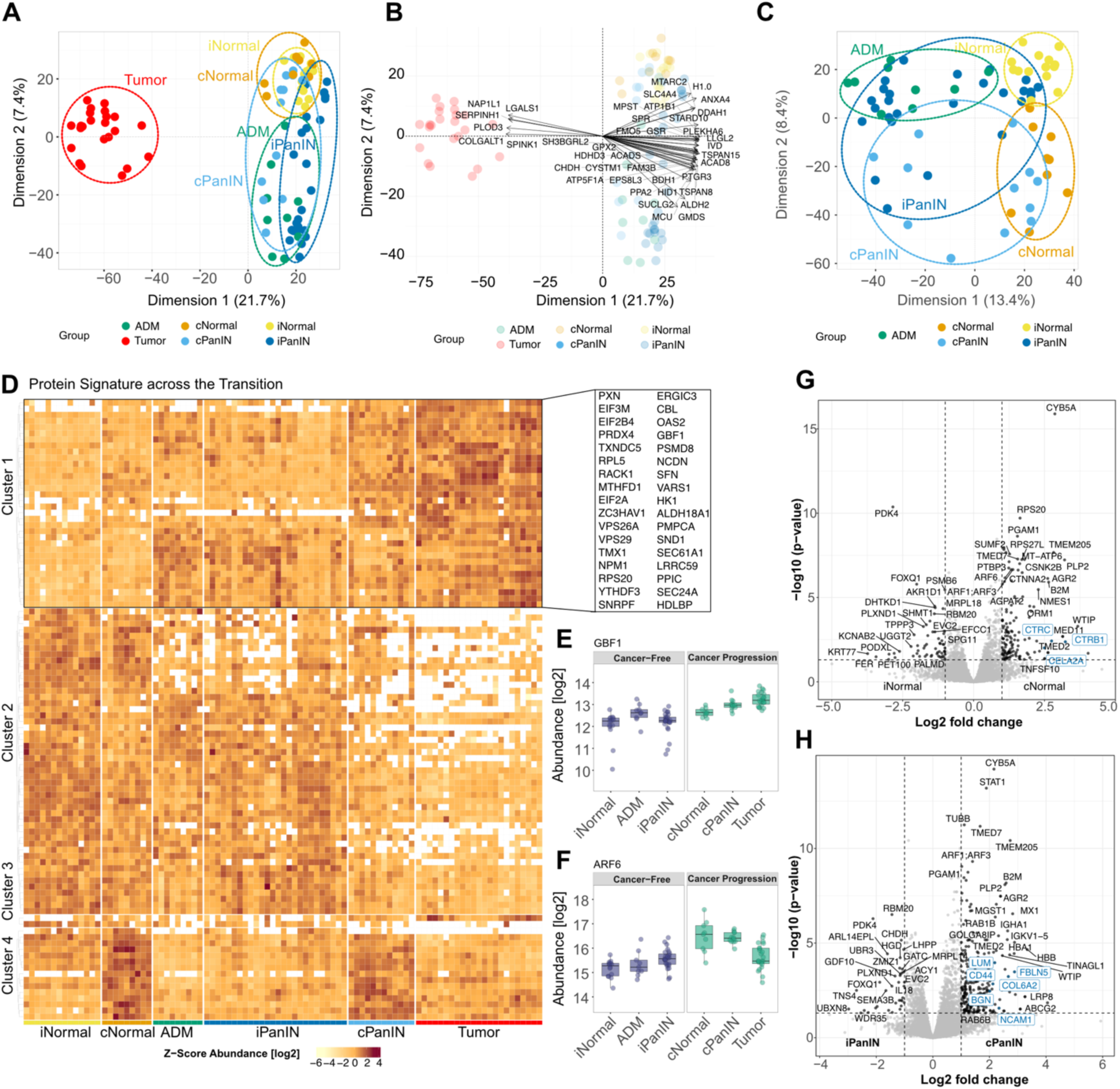
Protein signatures associated with PDAC tumorigenesis. **A)** Principal Component Analysis (PCA) of all cohort groups including Normal, ADM, PanIN, and tumor; dimensions of highest variance are shown (Dim.1 and Dim.2). **B)** Major drivers of data variance for the PCA analysis depicted in **(A)**. **C)** PCA of Normal, ADM, and PanIN; dimensions of highest variance are shown (Dim.1 and Dim.2). **D)** Signatures of normal-to-PanIN-to-PDAC tumorigenesis. ANOVA significance analysis and Post-Tukey extraction of proteins with significant differences in PDAC tumor vs. normal tissue while having consistent abundance levels between PDAC tumors and cPanINs. Protein names of Cluster 1 are highlighted (right). **E,F)** Representative boxplots of two clusters: GBF1 (Cluster 1) **(E)** and ARF6 (Cluster 4) **(F)**. **G)** Differential protein expression of normal tissue in the incidental and cancer-associated cohort (iNormal vs. cNormal). Significantly dysregulated proteins (fold change > 2; q-value < 0.05) are highlighted in black. Three digestive enzyme precursors are highlighted in blue. **H)** Differential protein expression of PanIN lesions in the incidental and cancer-associated cohort (iPanIN vs. cPanIN). Significantly regulated proteins (fold change > 2; q-value < 0.05) are highlighted in black. Proteins associated with ECM remodeling and adhesion are highlighted in blue.

### Deep Visual Proteomics reveals visually hidden molecular reprogramming

Focusing on non-tumor lesions, Principal Component Analysis (PCA) revealed that lesion morphology and disease association were the dominant factors shaping the molecular landscape of pancreatic tissue (**Fig. 2C**). The primary axis of variation (Dimension 1, accounting for 13.4% of total variance) separated histological classification of cells (Normal, ADM, and PanIN), while the secondary axis (Dimension 2, accounting for 8.4% of variance) stratified cancer-associated samples (“cCohort”) from cancer-free samples (“iCohort”). This analysis revealed that despite their morphological similarity, lesions from cancer-free and cancer-associated contexts exhibited profound molecular differences, underscoring how conventional histopathology can miss critical disease-relevant changes. Taken together, the principal components position “iNormal”, “cNormal”, “iPanIN”, and “cPanIN” along two axes that capture lesion type and cancer context (**Fig. 2C**). This two-dimensional distribution suggests that a subset of underlying molecular changes occur in parallel, rather than along a linear path. Overall, the ADM samples overlap with “iPanINs” but also form a distinct cluster, indicating that while they share some molecular features with established precursor lesions, they are not identical. Specifically, ADM samples form a separate cluster positioned between normal ducts and PanINs, providing the first molecular evidence in humans that they may represent an intermediate state in PDAC development, recapitulating findings observed in murine models.

Statistical analysis of protein expression across progression states revealed distinct molecular clusters with stage-specific dynamics (**Fig. 2D, Table S5**). For example, *Cluster 1* identified proteins demonstrating progressive upregulation from histologically normal epithelium and PanIN lesions to PDAC, collectively representing potential hallmarks of neoplastic commitment (**Fig. 2D, 2E**). In contrast, *Cluster 2* captured proteins whose expression peak in normal epithelium and PanINs, while attenuating in invasive lesions, underscoring the divergent molecular trajectories of protein networks occurring during early pancreatic tumorigenesis. Notably, *Cluster 4* revealed signatures of a cancer-associated “field effect”, comprised of proteins highly expressed in both “cNormal” and “cPanIN”, such as the vesicle trafficking component ARF6 (**Fig. 2D, 2F**), but absent in histologically comparable “iNormal” and “iPanINs”, respectively.

Based on the finding in *cluster 4*, we then conducted differential expression analysis by lesional subtype, which revealed profound molecular shifts driven by the cancer context, with markedly divergent protein expression profiles between cancer-free and cancer-associated samples from histologically identical regions (**Table S2**). Importantly, we collected cPanINs from tissue sections spatially separated from the tumor (different tissue blocks), eliminating the possibility that tumor cell contamination causes the observed molecular changes. The earliest detectable molecular event in PDAC tumorigenesis occurs in histologically normal ducts within the cancer-related microenvironment. Comparing “iNormal” with “cNormal”, we found significant molecular differences despite comparable morphological appearances. This molecular reprogramming of seemingly normal tissue aligns with genetic evidence of clonal spread through the ductal system, which has been shown to occur years before invasive carcinoma development (24). We identified 196 proteins with significantly altered abundance, with 70% (138 proteins) showing increased expression in cancer-associated “cNormal” ducts, indicating widespread cellular reprogramming involving epithelial remodeling and tumor microenvironmental influences (**Fig. 2G, Table S2**). For instance, “cNormal” ducts exhibited a coordinated loss of normal ductal identity, with markedly lower abundance of the structural keratin KRT77 and the adhesion regulator PODXL, but increased expression of physiological digestive enzyme precursors (CTRB1, CTRC, CELA2A) and immune surveillance components (TNFSF10/TRAIL, B2M) compared to the cancer-free “iNormal” state (**Fig. 2G**). Furthermore, we identified 298 significantly altered proteins between otherwise histologically identical grades (low-grade) of PanIN lesions, “cPanIN” *versus* “iPanIN” lesions, with the vast majority (88%, 262 proteins) showing increased abundance in “cPanIN”, highlighting the profound impact of the tumor-adjacent context on precursor lesion signatures (**Fig. 2H, Table S2**). These molecular patterns demonstrate that “cPanINs” possess a distinct molecular architecture associated with increased progression potential, characterized by enhanced stromal remodeling, immune modulation, metabolic adaptation, and pro-survival signaling compared to “iPanINs”. This was exemplified by the enrichment of proteins associated with ECM remodeling and cell adhesion (CD44, LUM, BGN, NCAM1, COL6A2, FBLN5), immune-related proteins (B2M, STAT1, IGHA1, IGKV1-5), vesicular trafficking (ARF1, RAB1B, RAB6B, TMED7), oxidative stress management (MGST1), glycolysis (PGAM1), and cell survival/resistance (LRP8, ABCG2, AGR2) (**Fig. 2H**). The distinct molecular profiles between “iPanINs” and “cPanINs” highlight that field cancerization drives molecular reprogramming even within otherwise histologically indistinguishable low-grade lesions.

### Molecular hallmarks of the cancer field effect and early PDAC evolution

Based on our comprehensive proteomic profiling, we delineated four critical molecular hallmarks that characterize the cancer field effect and progression to PDAC: 1) stress adaptation, 2) immune engagement, 3) metabolic reprogramming, 4) mitochondrial dysfunction (**Fig. 3A**). These coordinated transitions represent early molecular reprogramming events that precede overt histopathological changes, establishing potential windows for early detection and intervention.

**Fig. 3.**
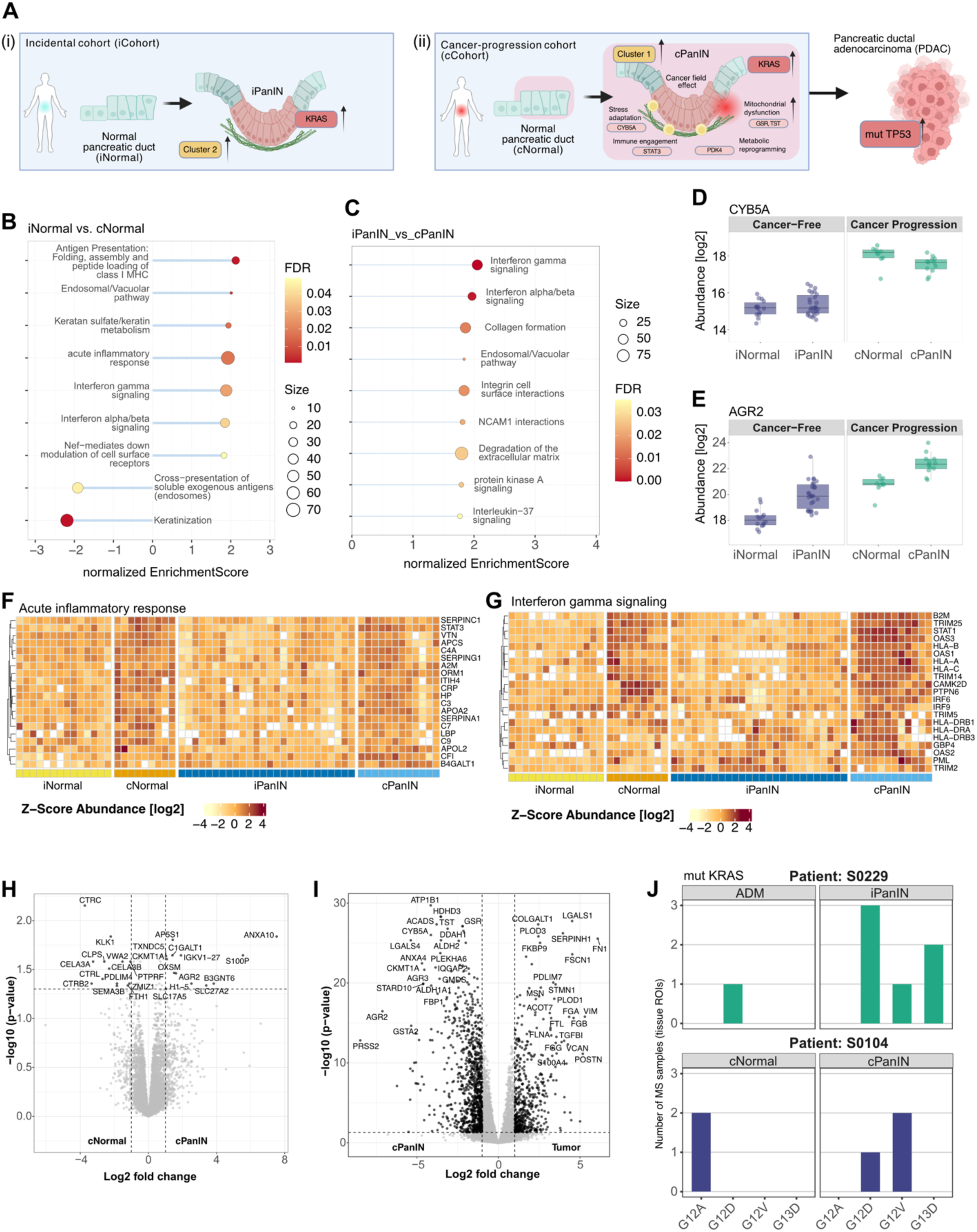
Cancer field effect in the proteomic reprogramming of normal-appearing pancreas. **A)** Schematic summary on pathways and distinct proteins involved in the multistep progression of human pancreatic neoplasia. Cluster 1 and 2 refer to the proteomic results shown in Fig. 2D. This figure was created using BioRender (https://www.biorender.com). **B,C)** Representative biological pathways enriched using GSEA comparing iNormal vs. cNormal **(B)** and iPanIN vs cPanIN **(C)**. FDR values are visualized by a color gradient while the number of proteins enriched for a pathway (size) is indicated by point size. **D,E)** Representative markers, CYB5A **(D)** and ARG2 **(E)**, associated with the cancer field effect in cancer-associated normal tissue and panIN lesion. **F,G)** Heatmap of z-score normalized protein abundances for the enriched pathways ‘Acute inflammatory response’ **(F)** and ‘Interferon gamma signaling’ **(G)**. **H,I)** Differential protein expression within the cancer-associated cohort comparing cNormal vs. cPanIN **(H)** and cPanIN vs. Tumor **(I)**. Significantly regulated proteins (fold change > 2; q-value < 0.05) are highlighted in black. **J)** Normalized peptide frequency of KRAS hotspot mutations, including G12A, G12D, G13D, and G12V, in normal ducts, ADM or PanIN lesions. One patient is shown exemplarily for each cohort (iCohort and cCohort), see also **Table S6.**

#### 1. Stress adaptation

Pathway enrichment analysis revealed significant upregulation of endosomal and vacuolar pathway signatures in “cNormal” ducts and “cPanINs” compared to their cancer-free counterparts (“iNormal” ducts and “iPanINs”, respectively) (**Fig. 3B, 3C**), suggesting activation of cellular stress response and adaptation programs in response to tumor-adjacent microenvironmental cues. Thus, “cNormal” ducts exhibited increased expression of stress-response proteins (WTIP, CYB5A, TAGLN) and proteins involved in ER function and trafficking (PLP2, TMED2, AGR2, MED11) compared to “iNormal” ducts (**Tables S2, S3**). Many of these markers and machinery also remained upregulated in “cPanINs” relative to “iPanINs” (**Tables S2, S3**), suggesting that early adaptive responses to tumor microenvironmental stress are sustained through pre-neoplastic progression. Notably, CYB5A, a protein involved in oxidative stress regulation and previously described as a tumor suppressor in PDAC (25), was upregulated 7.3-fold in “cNormal” compared to “iNormal”. It remained elevated, though at slightly lower levels, in “cPanINs” compared to “iPanINs” (**Fig. 3D, Table S2**). This expression pattern may reflect an early protective response aimed at maintaining redox homeostasis in the setting of inflammatory stress, potentially acting to constrain STAT3-mediated pro-inflammatory signaling imposed by the adjacent tumor (26). AGR2, a PDAC-associated oncogene that mediates a mucin-associated transcriptional program and ER stress responses (27–29) was upregulated 6.1-fold in “cNormal” compared to “iNormal” and showed a stepwise increase from “iNormal” to “cPanINs” (**Fig. 3E, Table S2**). This expression pattern provides compelling molecular evidence for cancer-associated alterations extending beyond histologically visible lesions.

#### 2. Immune engagement

Analysis of the cancer field effect revealed significant alterations in inflammatory signaling networks within histologically unremarkable ductal epithelium adjacent to cancer (“cNormal”). Pathway enrichment analysis demonstrated activation of inflammatory signaling networks in “cNormal” tissue and enhanced antigen presentation via MHC class I receptors, while presentation of soluble exogenous antigens pathways was downregulated (**Fig. 3B**). These immune alterations in “cNormal” ducts were accompanied by a pro-inflammatory state characterized by elevated acute-phase proteins (CRP, C3) and STAT3 compared to the incidental cohort (“iNormal”) (**Fig. S4A**). Detailed protein-level analysis of the acute inflammatory response revealed a progressive gradient of inflammatory activation across the neoplastic spectrum (**Fig. 3F**). Multiple acute-phase reactants and complement components showed coordinated upregulation, particularly in “cPanIN” lesions, including CRP, SERPINA1, SERPING1, complement components (C3, C4A, C7, C9), haptoglobin (HP), and alpha-2-macroglobulin (A2M). This inflammatory signature increased in intensity from “iNormal” to “cNormal” and then further from “iPanIN” to “cPanIN”, revealing a molecular state of inflammation before visible pathological changes appear. STAT3 upregulation correlated with the cellular response to inflammatory signals, consistent with a feed-forward signaling loop within the cancer field. The concurrent elevation of both CYB5A and STAT3 in both “cNormal” and “cPanINs” may represent an adaptive response to pro-inflammatory and stress signals within the tumor-adjacent, non-neoplastic milieu (22,30).

Interferon gamma signaling emerged as a dominant feature in the field effect, evidenced by coordinated upregulation of interferon-stimulated genes across the disease progression trajectory, peaking in “cPanIN” lesions (**Fig. 3G**). This signature included transcription factors (STAT1, IRF6, IRF9), MHC class I and II components (B2M, HLA-A/B/C, HLA-DR/DRA/DRB3), interferon-inducible GTPases (GBP4), viral defense enzymes (OAS1/2/3), and immune regulatory proteins (TRIM14/25, PML). Patterns of interferon response were accompanied by enhanced antigen presentation machinery, including upregulation of MHC class II components (HLA-DRA, CD74) (**Fig. S4B**) and a lysosomal enzyme (CTSB) (**Fig. S4C**). The concomitant increases in HMGB1 and SMAD3 further reflected innate immune activation and stromal reprogramming (**Fig. S4D**). The convergence of these pathways implies a potential last ditch “cancer avoidance” checkpoint at the “cPanIN” stage, where neoplastic clones are still restrained by the host immune response. These features would also align with the recent striking finding that shows widespread polyclonal “cPanIN” burden (estimated to number ∼1,000) in the cancer-adjacent pancreas (8) – it is likely that only one or a handful of clonally related PanIN lesions eventually progress to cancer while the rest are restrained by the prevalent host immune response. Intriguingly, the prominent upregulation of STAT1 and increased interferon signaling in “cPanINs” suggests a distinct emphasis on interferon-mediated immune responses in cancer-associated precursor lesions, but absent in “iPanINs” (**Fig. 3G, S4E**). Together, the data show that chronic inflammation, type I interferon signaling, antigen presentation, and plasma-cell infiltration co-emerge in the cancer field, establishing an interferon-driven microenvironment that likely shapes early progression.

#### 3. Metabolic reprogramming

Pancreatic precursor lesions are often characterized by increased glucose uptake and enhanced metabolic activity (31,32), however the timing of this reprogramming within the context of human precursor lesions remains unknown. We consistently observed markedly lower abundance of the regulatory enzyme pyruvate dehydrogenase kinase 4 (PDK4) in “cNormal” and “cPanIN” lesions compared to their cancer-free “iNormal” and “iPanIN” counterparts (**Fig. S4F**). Further, the glycolytic mediator phosphoglycerate mutase 1 (PGAM1) and PDK4 showed an inverse expression pattern between the “iCohort” and “cCohort”, respectively (**Fig. 2G, 2H**). Given that PGAM1 promotes glycolysis and PDK4 inhibits mitochondrial respiration and subsequent fatty acid metabolism through the pyruvate dehydrogenase (PDH) complex (33,34), this inverse relationship suggests a metabolic shift toward enhanced glycolytic metabolism and altered lipogenesis within the tumor-adjacent precursors, potentially supporting the increased biosynthetic demands of early neoplastic transformation.

The transition from “cNormal” to “cPanIN” also featured additional metabolic regulators (for example, the fatty acid transporter SLC27A2) (**Fig. 3H**), further reinforcing the central role of metabolic reprogramming in early preneoplastic transformation. Finally, glycosylation emerged as a consistent molecular marker of PanINs, accompanied by upregulation of glycosylation-related enzymes (B3GNT6, C1GALT1) (**Fig. S5**), underscoring the frequently observed glycosylation aberrations in PDAC (35,36).

#### 4. Mitochondrial dysfunction during PanIN transition to invasive cancer

The transition from “cPanIN” to invasive cancer represented the most substantial molecular shift observed in our study, marked by distinct alterations in cellular programs and protein abundance near tumor regions (**Fig. 3I**). Surprisingly, the transition involved coordinated downregulation of multiple mitochondrial proteins (TST, GSR, ACADS, CKMT1A), concurrent with upregulation of invasion-promoting factors such as FSCN1 (Fascin), STMN1 (Stathmin 1), and S100A4 (**Fig. 3I**). As noted previously, PGAM1 upregulation was already observed in “cPanIN” lesions (**Fig. 2H**), suggesting that glycolytic reprogramming precedes the mitochondrial dysfunction and facilitates the overt bioenergetic demands of an invasive phenotype. Of note, reduced expression of various mitochondrial membrane proteins (a measure of “mitochondrial fitness”) has been reported as a feature of tumor aggressiveness and metastatic propensity in PDAC (37). We therefore assessed whether these changes in mitochondrial proteins were associated with the classical and basal-like lineage states (38,39), as the latter is associated with an aggressive PDAC phenotype (40). Lineage marker analysis revealed that both “iPanIN” and “cPanIN” lesions predominantly retained a classical phenotype (elevated CLDN18, GATA6, HNF4A, **Fig. S6A**), while prototypical basal-like markers (S100A2, KRT17, HMGA2) progressively increased during malignant transformation (**Fig. S6B**). Basal-like PDAC cells express lower levels of mitochondrial pyruvate carrier 1 (MPC1) (41), and we observed ∼2-fold lower MPC1 expression in tumors compared to “cPanINs” (**Table S2**). Collectively, these findings suggest that glycolytic reprogramming and mitochondrial dysfunction are two of the metabolic perturbations that bridge the PanIN-PDAC transition and might be potential vulnerabilities that can be exploited for cancer interception.

### KRAS mutation-specific peptide profiling reveals polyclonality in pancreatic precursor lesions

Our spatial proteomics workflow detected multiple KRAS hotspot mutations as tryptic peptides in precursor lesions and histologically normal ducts, providing protein-level validation of pancreatic field cancerization (**Fig. 3J, Table S6**). Remarkably, we observed polyclonal KRAS mutations within individual patients - a finding that validates at the proteomic level previous genomic observations of multiclonal PanIN evolution (8).

In individual S0229 (a cancer-free organ donor), we detected the G12D mutation in an ADM lesion, while the corresponding iPanIN lesions harbored a diverse mutational spectrum including G12D, G12V, and G13D peptides (**Fig. 3J, top panel**). This polyclonality within a single cancer-free individual is the first demonstration that multiple independent KRAS-mutant clones can arise and coexist in the precursor landscape even outside of the context of an adjacent PDAC.

Even more striking, patient S0104 showed KRAS mutations in histologically normal ducts (”cNormal”), with rare G12A peptides detected (**Fig. 3J, bottom panel**). The same patient’s cPanIN lesions displayed additional mutations (G12D, G12V), revealing progressive clonal diversification from normal-appearing epithelium to established precursors. The presence of KRAS mutations in morphologically normal ducts provides direct proteomic evidence that molecular transformation precedes histological changes - a finding with profound implications for early detection strategies.

### Spatial proteomic dissection reveals functional niches and intra-tumoral heterogeneity in PDAC

Our multimodal foundation model framework integrated AI-driven tissue deconvolution with spatially resolved mass spectrometry-based proteomic mapping, revealing significant intra-tumoral heterogeneity in PDAC specimens (**Fig. 4**), recently described only at the transcriptional level (38). We performed multi-region proteomic sampling across three tumor cases with matched “cPanIN” lesions, demonstrating distinct molecular profiles between spatially separated regions within individual tumors (**Fig. 4A, 4C, 4E**). This spatial resolution exposed functionally diverse proteomic landscapes that would remain undetectable using conventional bulk tissue analysis. PCA demonstrated remarkable molecular divergence between spatial regions in all three cases (**Fig. 4B, 4D, 4F**), with certain tumor areas dominated by stromal remodeling programs, while others exhibited signatures associated with altered cellular physiology. In one case (S0115), certain tumor areas showed enrichment for ECM-remodeling proteins (FBLN2/5, FLNA, PXDN), cytoskeletal proteins (TNS1, PDLIMs), and immunoglobulins (IGHV4, IGKV3D.20, IGKV2.28), potentially associated with invasive behavior and immune responses (**Fig. 4B**). Adjacent regions within the same tumor exhibited elevated expression of ER stress-response and membrane dynamics proteins (ERP44, ESYT2, LPCAT3) (**Fig. 4B**). In these molecularly distinct tumor regions, cancer-related markers such as TP53 and DCLK1 exhibited divergent expression patterns (**Fig. S6C, S6D**). These spatially resolved molecular signatures uncover functional niches that conventional bulk analysis would obscure, indicating that regional adaptation may contribute to PDAC progression through coordinated programs of stromal remodeling and immune modulation.

**Fig. 4.**
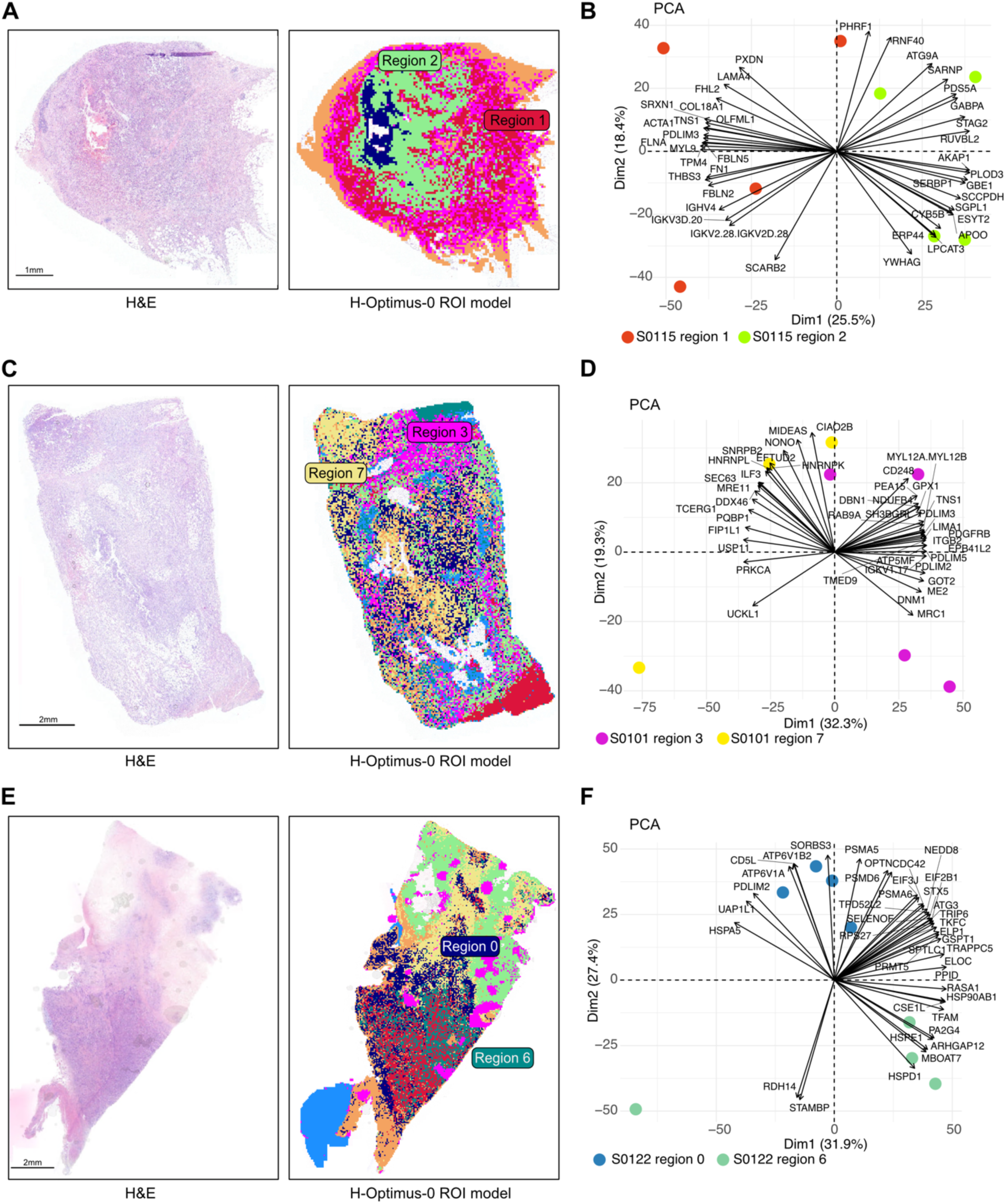
Resolving drivers of tumor heterogeneity and progression in mature PDAC. **A,C,E)** Computational pathology algorithms of this study resolving tumor heterogeneity in three PDAC tumors from patients S0115, S0101, and S0122. H&E staining is given as clinically standardized reference (left), based on which the AI model classified tumor heterogeneity (right). Regions 1 and 2 **(A)**, regions 3 and 7 **(C)**, and regions 0 and 6 **(E)** were identified by the pathology foundation model. \**B,D,F)** Principal component analysis (PCA) of proteomic data for three PDAC tumors, S0115 **(B)**, S0101 **(D)**, and S0122 **(F)**. Key drivers of variance are highlighted in black. The PCA shows distinct separation between the two regions, corresponding to regions of interest (ROI) annotated by a pathologist.

### The proteomic landscape of human acinar-to-ductal metaplasia

While murine acinar-to-ductal metaplasia (ADM) has been extensively profiled by single cell approaches (42,43), human ADMs – and their relationship to low grade PanIN lesions – remain largely unexplored. Our deep proteomics analysis in the earliest exocrine precursor lesions from cancer-free organ donors − ADM and “iPanIN” − revealed a molecular continuum with both shared and distinct features. Shared molecular programs define the precursor state. Compared with “iNormal”, both ADM and “iPanIN” lesions upregulate signatures of three core programs: oxidative stress response and metabolic adaptation (e.g. CKMT1A, PGC, OLFM4, and HK1), glycosylation and vesicle trafficking (e.g. GOLM1 and SYTL1), and cell adhesion and barrier integrity disruption (e.g. CLDN18, MUC1, MUC6, and S100P) (**Fig. 5A-5E**). Among these, CKMT1A (creatine kinase, mitochondrial 1A) emerged as the most highly upregulated protein in both ADM and “iPanIN” lesions, suggesting that altered energy metabolism may represent a hallmark of early ductal epithelial transformation. These shared features position ADM and “iPanIN” lesions along a common developmental trajectory, distinct from normal ducts. Moreover, direct comparison uncovered divergent programs between ADM and “iPanIN” lesions. ADM preferentially expressed metabolic reprogramming and ductal/progenitor markers (**Fig. 5F**), including PHGDH (serine synthesis, **Fig. 5G**) and PIKFYVE (lipid metabolism).

**Fig. 5.**
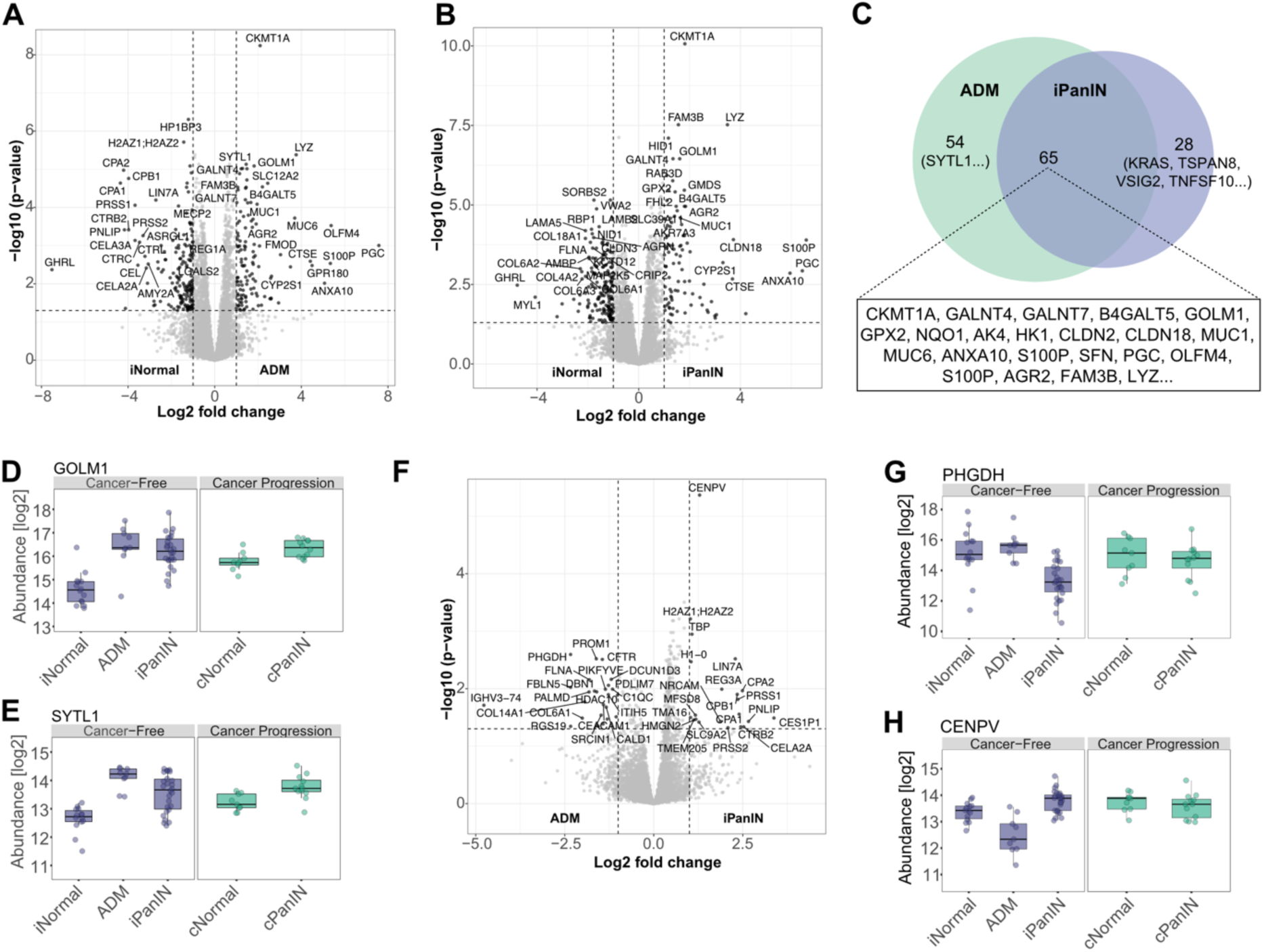
Early proteome changes in lesions of the cancer-free cohort. **A,B)** Differential protein expression of iNormal vs. ADM **(A)** and iNormal vs. iPanIN **(B)**. Significantly regulated proteins (fold change > 2; q-value < 0.05) are highlighted in black. **C)** Venn diagram showing unique and overlapping upregulated proteins in ADM or iPanIN compared to iNormal. Representative overlapping proteins are highlighted. **D,E)** Novel markers of the ADM: GOLM1 **(D)** and SYTL1 **(E)**. **F)** Differential protein expression of ADM vs. iPanIN **(A)**. Significantly regulated proteins (fold change > 2; q-value < 0.05) are highlighted in black. **G,H)** Representative proteins exhibiting divergent expression patterns between ADM and iPanIN: PHGDH **(G)** and CENPV **(H)**.

In contrast, iPanINs selectively upregulated chromatin regulation and epithelial architecture proteins: CENPV (chromatin remodeling, **Fig. 5H**) and LIN7A (cell polarity). This molecular divergence reveals ADM as a metabolically active transition state, while “iPanIN” lesions establish the structured epithelial program characteristic of stable precursor lesions.

## Discussion

Our AI-powered spatial proteomic atlas represents the first high-resolution protein-level molecular characterization of both incidental and cancer-associated precursors in the development of PDAC. Our work defines molecular changes across four critical transitions. Our central discovery - that histologically normal ducts from cancer patients display significantly different molecular profiles than those from cancer-free individuals despite identical histopathological presentation - represents an important advance with immediate implications for risk stratification and early intervention strategies.

Histologically normal pancreatic ducts associated with cancer exhibit extensive molecular alterations despite their normal appearance under conventional microscopy. We identified a distinctive molecular signature of field cancerization characterized by stress adaptation, immune engagement, metabolic reprogramming, and mitochondrial dysfunction. The striking upregulation of the established oncogene AGR2 (6.1-fold) in morphologically normal ducts provides compelling evidence that field cancerization extends substantially beyond visible lesions. The marked increase in CYB5A expression (7.3-fold) suggests it may constrain the pro-tumorigenic environment within the non-tumor milieu. This molecular field effect helps explain the high recurrence rates following surgical resection, as residual tissue that appears histologically normal may harbor pro-tumorigenic molecular environments.

Our findings align with recent spatial transcriptomic analyses by Bell et al. (9) and Carpenter et al. (6), who documented transcriptional reprogramming in the PDAC microenvironment. However, our proteomic approach extends these observations by revealing actionable protein-level alterations with higher resolution between tissue compartments, particularly demonstrating more pronounced molecular differences between “iPanIN” and “cPanIN” lesions than reported at the transcriptome level. While Carpenter et al. concluded that similarities far outweighed any differences between sporadic and tumor-associated PanINs at the RNA level (6), our proteomic data reveals substantial molecular divergence between these histologically similar lesions. The coordinated inflammatory signatures in cancer-associated tissues raise a crucial question: Does inflammation drive neoplastic transformation rather than merely responding to it? Our data show coordinated activation of STAT signaling pathways – including prominent upregulation of both STAT3 and STAT1 in “cPanIN” lesions – predicating an emphasis on interferon-mediated immune responses within the soil on which cancer arises. Together with the enrichment of interferon signaling and acute inflammatory response programs in cancer-associated normal ducts, our findings suggest that inflammation actively drives rather than follows transformation. This pattern likely facilitates inflammatory remodeling of the early tumor-promoting immune microenvironment which has been shown to accelerate KRAS-driven carcinogenesis (10).

Our mass spectrometry-based detection of polyclonal KRAS mutations provides protein-level validation of recent genomic findings (8), although it extends those findings into cancer-free organ donor samples. Detecting multiple distinct KRAS mutations within individual patients - including in histologically normal ducts - demonstrates that molecular transformation precedes histological changes probably by years or decades. This has immediate implications for surveillance strategies, as the detection of shed mutant KRAS alleles within proximal biospecimens like duodenal juice (and less likely, in the peripheral circulation) has the potential to result in significant false positives. The detection of the rare G12A mutation in normal ducts is particularly intriguing. While G12D and G12V represent the most common KRAS mutations in established PDAC, G12A comprises less than 1% of cases (44). Its enrichment in our earliest lesions suggests that the mutational spectrum in precursor states may differ from that observed in invasive carcinoma, potentially reflecting differential selection pressures or varying oncogenic potencies among KRAS alleles.

These discoveries enable new strategies for early detection. The field effect molecular signature (CYB5A, WTIP, B2M, AGR2) could identify high-risk field cancerization before visible lesions appear, potentially through analysis of pancreatic fluid or liquid biopsies. Genetic studies estimate a median latency period of several years from the common ancestor to clinically apparent cancer (24), presenting a window to identify at-risk patients far earlier than current imaging-based approaches permit. The distinct molecular profiles of “cPanIN” versus “iPanIN” lesions provide criteria for risk stratification beyond histological grading. Markers such as STAT1, CD44, and interferon signaling components could distinguish high-risk from low-risk PanINs, informing surveillance strategies and intervention timing.

Despite our unprecedented insights provided by our study, several limitations warrant consideration. Our analysis focused primarily on empirical protein-level alterations without assessing corresponding epigenetic modifications and transcriptomic profiles that could provide additional mechanistic insight. Future work should validate these findings in larger cohorts, establish causal relationships between molecular alterations and disease progression, develop clinical assays for promising biomarkers, and extend this approach to other precursor lesions such as intraductal papillary mucinous neoplasms (IPMNs). Functional studies examining the consequences of manipulating key proteins identified in our analysis could further elucidate their roles in PDAC progression. The proteomic depth of this study significantly exceeds previous datasets in pancreatic precursor lesions (45), which were typically been limited by sample quantity, technological constraints, and reliance on mouse models rather than human tissue, enabling unprecedented insights into disease mechanisms.

In conclusion, our spatial proteomics platform unveils a novel molecular landscape of PDAC precursors and significantly enhances our understanding of neoplastic transformation trajectories. Molecular reprogramming at the protein level precedes histological changes, establishing a new paradigm for risk assessment and early intervention. By providing the proteomic counterpart to established genetic models of clonal evolution and spread, our work completes a more comprehensive picture of PDAC development. The four molecular transitions we identified - from field effect to mitochondrial dysfunction - provide a framework for developing stage-specific biomarkers and therapeutic targeting strategies. This approach offers the potential to transform PDAC clinical management from late-stage treatment to early interception, when curative intervention remains possible.

## Methods

### Study design and cohort

The study cohort included tissue specimens from 10 cancer-free organ donors and 5 treatment-naïve patients with PDAC. Tissues were obtained post-mortem within 4 hours through the University of Nebraska Medical Center (UNMC) Normal Organ Recovery Program (NORs) and Rapid Autopsy Program (NRAP). Non-cancer organ donor tissues were obtained through cooperation with Live On Nebraska. Formalin-fixed paraffin-embedded (FFPE) tissue blocks were sectioned at 3μm thickness and mounted on pre-coated 2.0 mm PEN membrane slides (MicroDissect GmbH) at MD Anderson Cancer Center (Houston, TX). A total of 19 tissue specimens (11 from organ donors including 2 samples from the same donor and 8 from PDAC patients) were selected based on the availability and quality of FFPE blocks suitable for laser microdissection.

All lesions were reviewed by A.M., an expert in pancreatic pathology with over two decades of expertise in PDAC and PanINs (46–48) as “ground truth” for histological classification. In organ donor pancreas, normal duct epithelium (iNormal), acinar-to-ductal metaplasia (ADM), and incidental PanINs (iPanINs) were identified. In three of five PDAC patients, non-tumor tissue blocks adjacent to invasive carcinoma − containing cancer-associated PanINs (cPanINs) and corresponding normal ducts (cNormal) − were included in the analysis. Tumor regions were analyzed from all five PDAC cases. We note that cPanINs and corresponding tumor samples were collected from different tissue sections on separate blocks. This spatial separation ensures that the molecular signatures identified in cPanINs represent true field effects rather than contamination from adjacent tumor cells.

All procedures were conducted in accordance with the Declaration of Helsinki and approved by the UNMC Institutional Review Board (IRB 091-01), with written informed consent obtained prior to enrollment. All patient and donor data were de-identified and anonymized. Cohort metadata, including age and gender, are provided in **Table S7**.

### AI image analysis

Tissue specimens were prepared and stained with H&E and imaged as described in the **Method Supplement**. Each acquired whole-slide image (WSI) was stored in ‘.mrxs’ format and processed using a custom tile extraction pipeline. Initially, each WSI was down-sampled to a lower resolution to enable tissue-background segmentation. The resulting tissue regions were partitioned into non-overlapping tiles of 224 × 224 pixels at a resolution of 0.24 microns per pixel (mpp), corresponding to a physical tile size of 54 × 54μm. Across 21 slides, a total of 1,618,248 tiles were extracted, resulting in an average of 77,059 tiles per slide. The tile resolution of 0.24mpp was chosen based on initial experiments showing it effectively captures relevant tissue morphologies. The extracted tiles were passed to H-optimus-0 (49), a vision transformer-based foundation model for pathology that consists of 1.1 billion parameters and was pretrained on over 500,000 WSIs. Each tile was embedded into a 1,536-dimensional feature vector. Model inference was executed on a local workstation equipped with 2 x 24GB NVIDIA GeForce RTX 4090 GPUs. Subsequently, k-means clustering was applied to the feature vectors of all tiles across all slides from a cohort, with the number of clusters *k* varied in the range *k* ∈ {2,…, 50}. This procedure is referred to as global clustering. The number of clusters *k* was adjusted interactively based on expert input on a subset of slides of the sample cohort. Based on this input, the clustering configuration was set to *k* = 23 for the iCohort and *k* = 10 for the cCohort and applied to all remaining slides. For the PDAC cohort each slide was clustered independently using k-means, with the number of clusters *k* varied within the range *k* ∈ {2,… 10}. This local clustering approach enables the identification of intra-slide tumor heterogeneity, in contrast to detecting morphological patterns shared across the entire cohort. All slides were subjected to human pathology review. Throughout the manuscript, we refer to each cluster as ‘region’.

### Deep Visual Proteomics

Details on the multi-modal analysis pipeline are provided in the **Method Supplement**. In brief, the vision transformer-based foundation model for AI-image processing guided a laser microscope developed for the exact requirements of this pipeline (LMD7, Leica Microsystems). 100 phenotypically matched cells were extracted by gravity-pulsed laser collection into a 384-well plate (Eppendorf twin.tec® PCR Plate 384 LoBind®, 0030129547). The resulting samples were prepared on an automated liquid handling platform (Agilent Bravo) for mass spectrometry, streamlining cell lysis and enzymatic digestion of proteins into peptides using trypsin and LysC. For ultra-sensitive LC-MS/MS, peptide samples were analyzed on an Orbitrap Astral mass spectrometer (Thermo Fisher Scientific) coupled to an Evosep One liquid chromatography System (Evosep Biosystems) using a 40 samples-per-day (40 SPD) Whisper-zoom standardized gradient (32.5 min). Resulting MS data were processed using the DIA-NN software suite (v1.8.1).

### Data analysis and statistics

All analyses were performed in the R statistical environment (v4.3.2). Protein raw intensities were log2 normalized and MS measurement grouped biologically. Protein IDs were determined based on the ‘stats.tsv’ output file; ‘pg_matrix.tsv’ was used for all statistical and data analyses. Plots were generated using the ggplot2 package (v3.5.0). Microscopy images were exported from QuPath (v0.5.1). Figures were generated in the ‘Affinity Designer’ licensed software and schematics in BioRender. Details on the data statistics are given in the **Method Supplement**.

### Search of KRAS Variants

A custom FASTA file was generated containing the human KRAS protein sequence with clinically relevant cancer-associated variants. The following amino acid substitutions were included: G12A, G12C, G12D, G12R, G12V, G13C, G13D, Q61H, Q61R, and A146T. These variants represent the most frequently observed mutations in KRAS-driven cancers. Theoretical spectral libraries were generated using AlphaPeptDeep version 1.3.0. The KRAS variant FASTA file and a default FASTA w/o KRAS were processed with the following parameters: trypsin digestion allowing up to 2 missed cleavages, peptide charge states from +2 to +3, and normalized collision energy (NCE) of 25 with the ‘Lumos’-setting. Variable modifications included phosphorylation on serine, threonine, and tyrosine residues (Phospho@S/T/Y), acetylation on lysine (Acetyl@K), and ubiquitin remnant on lysine (GlyGly@K). Carbamidomethylation of cysteine (Carbamidomethyl@C) was set as a fixed modification. The predicted spectral libraries were merged and subsequently used to search data using DIA-NN version 1.8.1 with default settings.

## Data availability

Pseudonymized MS-based proteomics data of this study have been deposited to the ProteomeXchange Consortium via the PRIDE partner repository. These data will be publicly available upon the publication of this work.

## Acknowledgements

Tissues used in this study were obtained through the UNMC Rapid Autopsy Program and Normal Organ Recovery Program, in cooperation with Live On Nebraska. We gratefully acknowledge and thank the donors, patients, and their families for their invaluable contributions to scientific research. This work was funded by OmicVision Biosciences. A. Maitra was supported by the Sheikh Khalifa bin Zayed Foundation, U54CA274371, U01CA200468 and U24CA274274. P. M. Grandgenett was supported by U01CA210240, P30CA36727, and R50CA211462. J. Min was supported by a Pilot/Feasibility Award from the Texas Medical Center Digestive Disease Center (P30DK056338) and a Seed Grant from the Hirshberg Foundation for Pancreatic Cancer Research. We thank Constantin Ammar for the constructive discussion regarding the KRAS variant search.

## Authors’ Contributions

**A. J. Min**: Conceptualization, Data curation, Investigation, Visualization, Writing-original draft, Writing-review and editing. **L. Schweizer**: Conceptualization, Data curation, Formal analysis, Investigation, Visualization, Writing-review and editing. **G. Zonderland**: Data curation. **B. C. Selvanesan**: Data curation. **L. Oldenburg**: Formal analysis, Methodology. **S. Bae**: Data curation. **B. J. Kim**: Data curation. **B. J. Swanson**: Resources. **K. A. Klute**: Resources. **T. C. Caffrey**: Resources. **P. M. Grandgenett**: Conceptualization, Resources, Writing-review and editing. **M. A. Hollingsworth**: Conceptualization, Resources. **I. Ummat**: Funding acquisition, Writing-review and editing. **M. T. Strauss**: Formal analysis, Methodology. **A. Mund**: Conceptualization, Supervision, Project administration, Writing-original draft, Writing-review and editing. **A. Maitra**: Conceptualization, Supervision, Project administration, Writing-review and editing.

