## Supplementary data for "AI-powered Deep Visual Proteomics reveals critical molecular transitions in pancreatic cancer precursors"

[#]Corresponding authors

### Supplementary Data

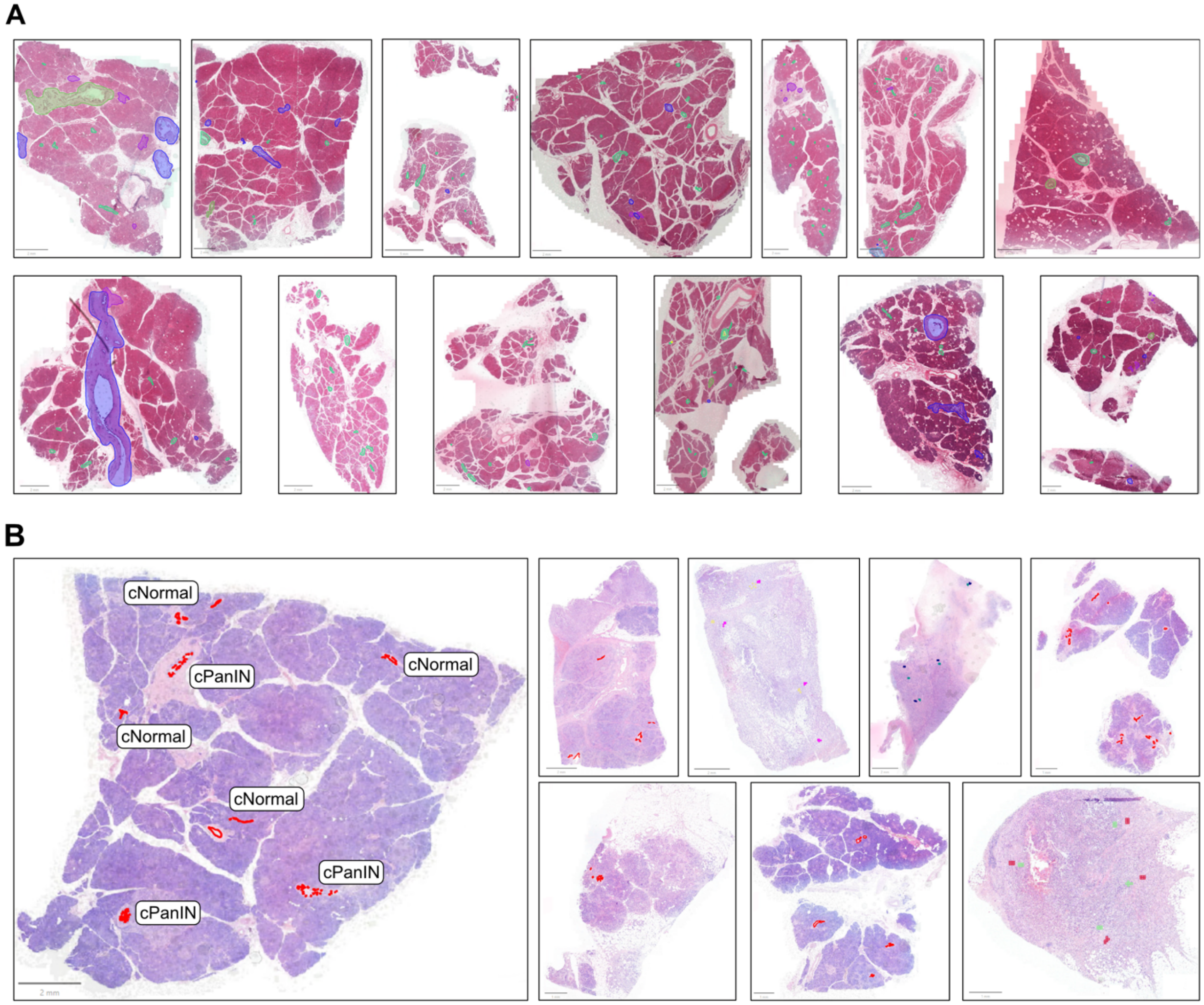


**Fig. S1 | Cohort overview.**

**A,B)** Whole slide images of H&E-stained tissue specimen for the incidental **(A)** and cancer-associated **(B)** cohorts. Regions of interest (ROI) are highlighted and annotated exemplarily for one slide.

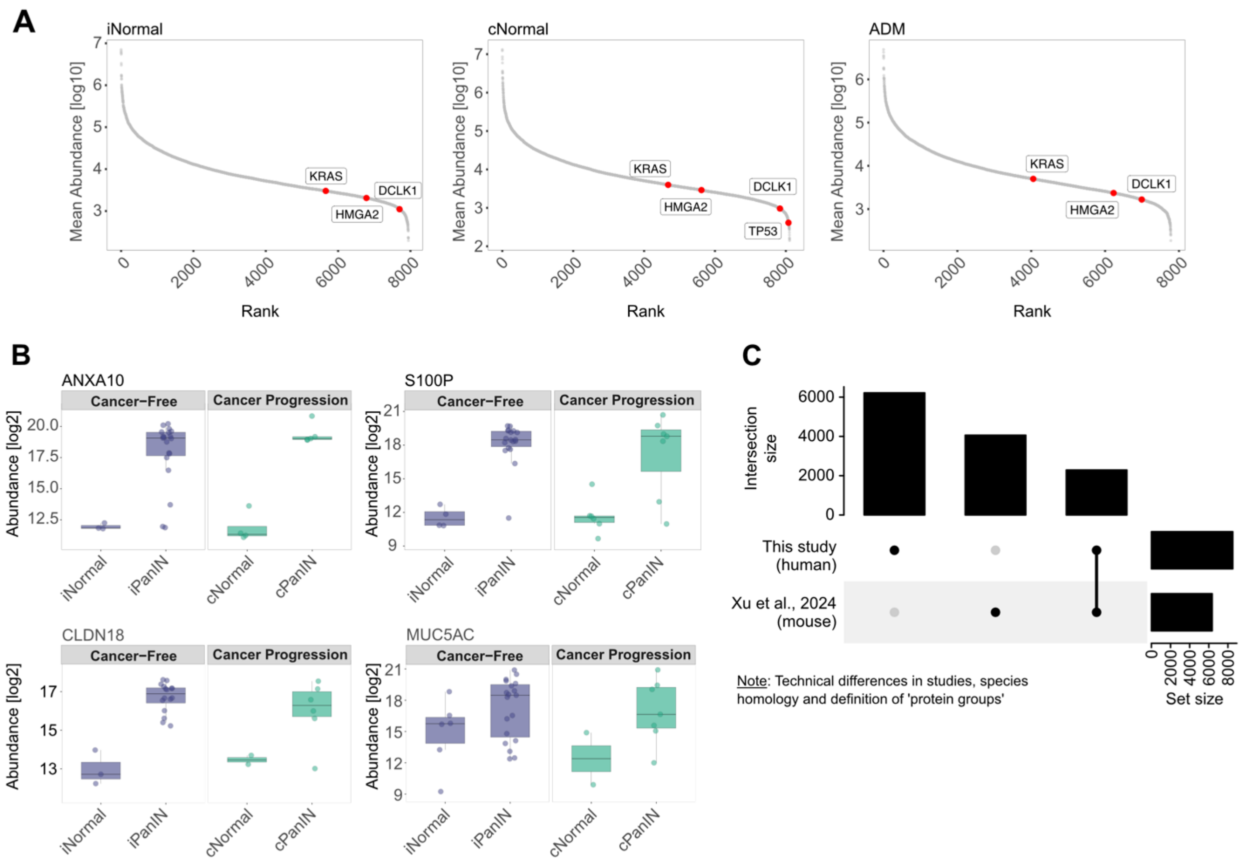


**Fig. S2 | Study quality control – reference work.**

**A)** Mean ranked abundance of proteins found in iNormal, cNormal, and ADM groups. The abundance of key markers is highlighted in red (KRAS, HMGA2 DCLK1, TP53).

**B)** Expression of canonical PanIN markers, ANXA10, S100P, CLDN18, and MUC5AC, across iNormal, iPanIN, cNormal, and cPanIN groups.

**C)** Upset plot comparing this study to recent work of Xu et al (45). Note the technical differences of the studies based on species (human vs. mouse) and definition of ‘protein groups’.

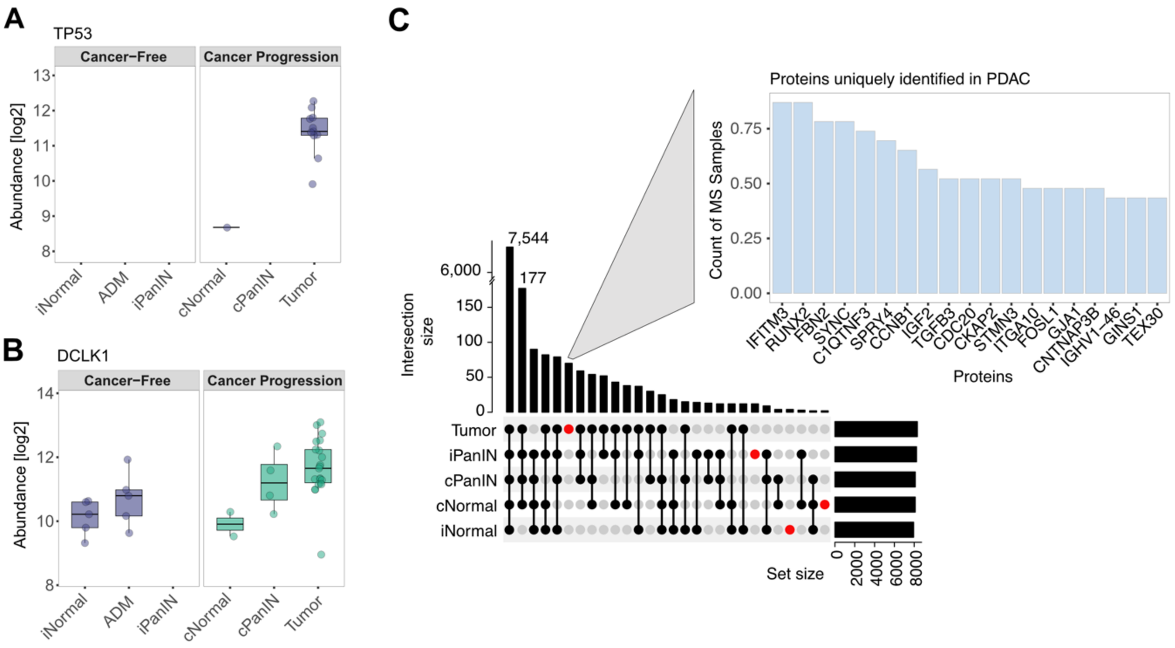


**Fig. S3 | Distinct protein abundance patterns across PDAC progression.**

**A,B)** Protein abundance for TP53 **(A)** and DCLK1 **(B)** summarized across cohorts.

**C)** Upset plot identifying proteins with unique abundance in a specific group. Of note, IFITM3 and RUNX2 were expressed uniquely in more than 97% of all PDAC tumor samples, while being entirely absent in normal or PanIN lesions.

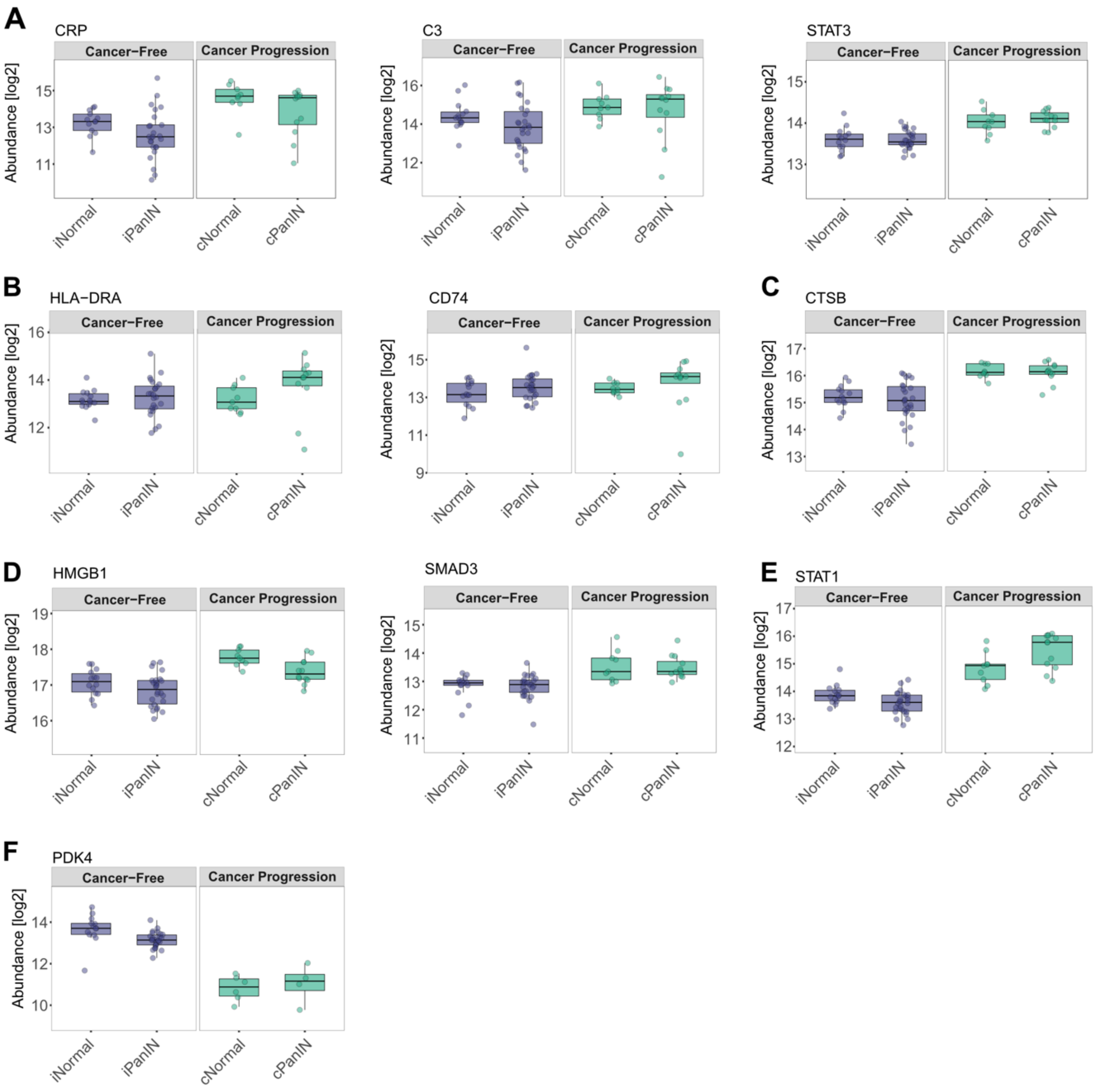


**Fig. S4 | Detailed abundances on proteins involved in the cancer field effect.**

Boxplots showing elevated levels of acute-phase proteins **(A)**, MHC class II components **(B)**, a lysosomal enzyme **(C)**, markers of innate immune activation and stromal reprogramming **(D)**, the inflammatory signaling-related protein STAT1 **(E)**, and the metabolic regulator PDK4 **(F)**.

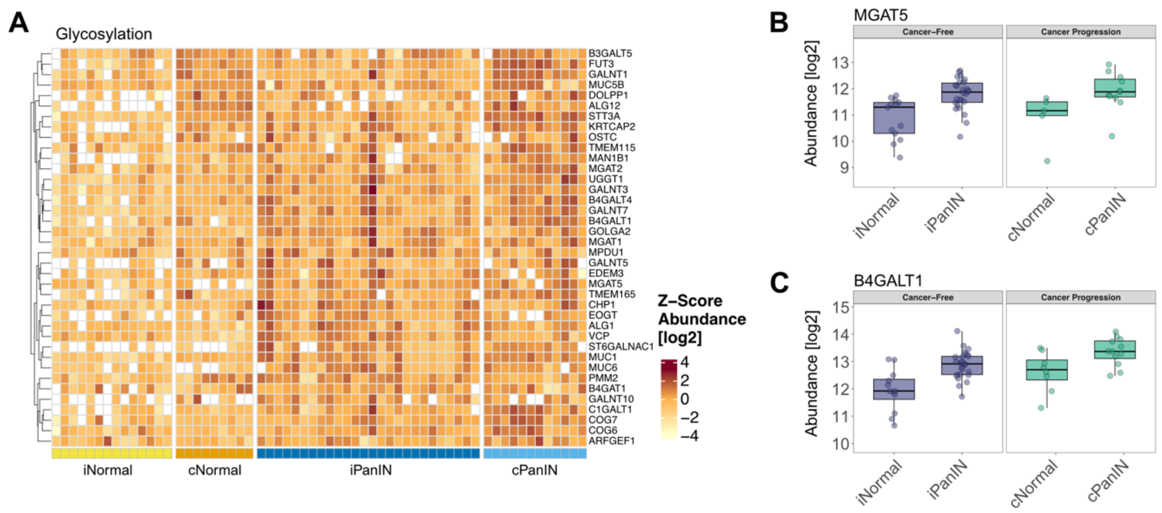


**Fig. S5 | Metabolic reprogramming in PanIN lesions.**

**A)** Heatmap of z-score normalized protein abundances for the enriched pathway 'Glycosylation'.

**B,C)** Representative glycosylation-related proteins, MGAT5 **(B)** and B4GALT1 **(C)**, identified as potential indicators of early PDAC precursor lesions.

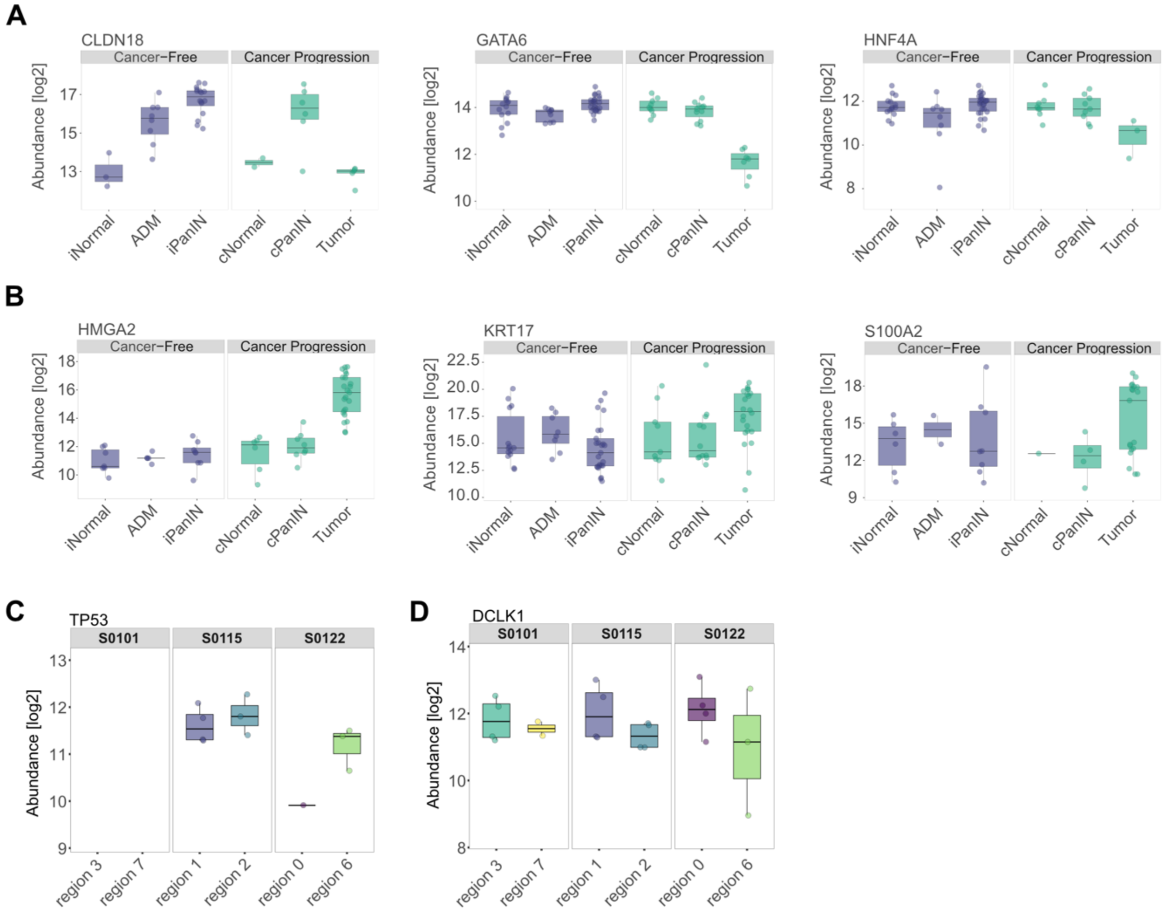


**Fig. S6 | Characteristic PDAC tumor markers in the investigated trajectory.**

**A,B)** Protein markers of the classic PDAC phenotype **(A)** and the more aggressive variant showing basal-like cell markers **(B)**.

**C,D)** Protein abundance for TP53 **(C)** and DCLK1 **(D)** across distinct tumor regions in individual patients. Double-pseudonymized patient IDs are shown.

**List of supplementary tables**

Table S1: Deep Visual Proteomics data

Table S2: Differential expression

Table S3: Biological pathway enrichment (GSEA)

Table S4: Unique identifications via mass spectrometry

Table S5: ANOVA heatmap data

Table S6: KRAS peptide mutations

Table S7: Patient demographics
